# Mixture distributions in a stochastic gene expression model with delayed feedback

**DOI:** 10.1101/855783

**Authors:** Pavol Bokes, Alessandro Borri, Pasquale Palumbo, Abhyudai Singh

## Abstract

Noise in gene expression can be substantively affected by the presence of production delay. Here we consider a mathematical model with bursty production of protein, a one-step production delay (the passage of which activates the protein), and feedback in the frequency of bursts. We specifically focus on examining the steady-state behaviour of the model in the slow-activation (i.e. large-delay) regime. Using a quasi-steady-state (QSS) approximation, we derive an autonomous ordinary differential equation for the inactive protein that applies in the slow-activation regime. If the differential equation is monostable, the steady-state distribution of the inactive (active) protein is approximated by a single Gaussian (Poisson) mode located at the globally stable steady state of the differential equation. If the differential equation is bistable (due to cooperative positive feedback), the steady-state distribution of the inactive (active) protein is approximated by a mixture of Gaussian (Poisson) modes located at the stable steady states; the weights of the modes are determined from a WKB approximation to the stationary distribution. The asymptotic results are compared to numerical solutions of the chemical master equation.

## 1 Introduction

Gene expression in individual cells involves the interaction of molecules which are present at low copy numbers (Eldar and Elowitz, 2010; Munsky et al, 2012). The intrinsic noise generated by the low-copy-number reactions is passed down to the end product of gene expression, the protein, and results in temporal fluctuations and cell-to-cell heterogeneity of the protein copy number (Taniguchi et al, 2010; Suter et al, 2011). The production of proteins in bursts of multiple copies is one of the most important sources of protein variability (Singh et al, 2010; Dar et al, 2012). Specifically, bursty production accounts for super-Poissonian variability observed in protein copy numbers (Thattai and van Oudenaarden, 2001).

Mathematical modelling has been proved useful in understanding the mechanisms of stochastic gene expression. The underlying probability distributions are typically defined as solutions to a specific master equation (Paulsson, 2005; Veerman et al, 2018; Albert, 2019). Explicit solutions to the master equation, especially at steady state, can be found for models with few components (Bokes et al, 2012; Zhou and Liu, 2015) and/or with special structural properties (Kumar et al, 2015; Anderson and Cotter, 2016). Generally, however, explicit solutions are unavailable or intractable and one resorts to stochastic simulation or seeks a numerical solution to a finite truncation of the master equation (Munsky and Khammash, 2006; Borri et al, 2016; Gupta et al, 2017). An alternative approach, which often provides useful qualitative insights into the model behaviour, is based on reduction techniques such as quasi-steady-state (Srivastava et al, 2011; Kim et al, 2014) and adiabatic reductions (Bruna et al, 2014; Popovic et al, 2016), piecewise-deterministic framework (Lin and Doering, 2016; Lin and Buchler, 2018), linear-noise approximation (Schnoerr et al, 2017; Modi et al, 2018), or moment closure (Singh and Hespanha, 2007; Andreychenko et al, 2017; Gast et al, 2019).

Production delay is an inevitable part of gene expression (Monk, 2003; Zavala and Marquez-Lago, 2014; Bokes et al, 2018). It can be caused by a number of mechanisms, e.g. transcriptional/translational elongation (Roussel and Zhu, 2006), post-translational modification (Gedeon and Bokes, 2012), or compartmental transport (Mor et al, 2010; Sturrock et al, 2017). The delay specifies the amount of time that needs to pass before a newly produced molecule can partake in its regulatory function (specifically in feedback). Delay can be fixed or randomly chosen from a distribution (Barrio et al, 2006; Lafuerza and Toral, 2011; Gupta et al, 2014). Exponentially distributed delays are the simplest among distributed delays as they are realised by the passage of a single memoryless step. Erlang and phase-type distributions provide a wider family of distributed delays which can be generated by a finite network of memoryless states (Soltani et al, 2016). Previous results indicate that large one-step (exponential) and multi-step (Erlang/phase-type) delays reduce the super-Poissonian noise in a bursty protein down to Poissonian levels (Singh and Bokes, 2012; Stoeger et al, 2016; Smith and Singh, 2019). This effect is also seen experimentally with buffered noise in cytoplasmic mRNA levels compared to nuclear mRNA levels due to transport delays (Battich et al, 2015). Additional effects of the inclusion of a delay are observed if the protein regulates, via transcriptional feedback, its burst frequency. In case of negative feedback, delays of moderate size lead to an increase, rather than decrease in protein noise (Smith and Singh, 2019). In case of non-cooperative positive feedback, noise-driven bimodal protein distributions, which are observed in the absence of delay, turn unimodal upon the inclusion of a distributed delay, and eventually converge to the Poissonian statistics as the delay increases (Borri et al, 2019). In case of cooperative positive feedback, the introduction of delay has been reported to enhance the stability of the modes of the protein distribution (Gupta et al, 2013; Feng et al, 2016; Kyrychko and Schwartz, 2018).

In the paper we focus, for its relative simplicity, on the case of exponential delay. We refer to the delay as activation and distinguish between the inactive and active protein species. We will argue that in the limit of slow activation rates the model behaviour becomes deterministic at the level of the inactive protein. Behaviour of stochastic models near a deterministic limit can be interpreted using the large-deviation theory (Tsimring and Pikovsky, 2001; Heymann and Vanden-Eijnden, 2008; Kumar and Kulkarni, 2019) and quantified by WKB asymptotic approximations (Schuss, 2009; Bressloff, 2014; Assaf and Meerson, 2017). The WKB approach has been successfully applied to stochastic reaction kinetics systems with large molecule copy numbers (Hinch and Chapman, 2005; Be’er and Assaf, 2016; Yin and Wen, 2019), fast switching of internal states (Newby, 2012; Lin and Galla, 2016), or a combination of both (Newby and Chapman, 2014, discussed at greater length in Section 7). Here we will use the WKB approximation to obtain reliable estimates of the stationary distribution of the active (and also the inactive) protein in the slow activation (large-delay) regime.

The outline of the paper is as follows. The stochastic model is introduced in Section 2 and reduced in Section 3 to a deterministic rate equation by means of a quasi-steady-state (QSS) reduction. Section 4 introduces the WKB solution and outlines the key aspects of the WKB analysis. Section 5 compares the WKB and numerical solutions to the master equation. The WKB analysis is performed in Section 6. Section 7 concludes the paper with a discussion.

## 2 Model formulation

The paper is concerned with a reaction system involving an inactive protein X and an active protein S which are subject to the production, activation, and decay reaction channels (Table 1). Each reaction is specified by its rate and reset map. The reaction rate, upon the multiplication by the length of an infinitesimally short time interval, gives the probability that the reaction will occur within the interval. The reset map determines the adjustment of the reaction species copy numbers resulting from an occurrence of the reaction.

**Table 1:** Reaction channels in the delayed feedback model. The copy number of the inactive protein X is denoted by *X* (in italic type) and that of the active protein S by *s*. The production burst rate *f*_*s*_ is a function of *s*, which implements the feedback. The production burst size *B* is in general drawn from a prescribed random distribution. The parameter *ε* ≪ 1 determines the discrepancy between the *O*(1) slow activation timescale and the *O*(*ε*) fast timescale of turnover of the active protein S. The reset map of the activation channel ensures that *s* never exceeds an upper bound *s*_max_.

| Name | Scheme | Rate | Reset map |
| --- | --- | --- | --- |
| Production | $\emptyset \rightarrow B \times X$ | $\varepsilon^{-1} f_s$ | $X \rightarrow X + B$ |
| Activation | $X \rightarrow S$ | $X$ | $X \rightarrow X - 1,$<br>$s \rightarrow \min\{s + 1, s_{\max}\}$ |
| Decay | $S \rightarrow \emptyset$ | $\varepsilon^{-1} s$ | $s \rightarrow s - 1$ |

We are specifically interested in studying the model in the regime of slow activation. Making activation slow is equivalent to making all the remaining reactions fast: indeed, by Table 1, the activation rate is *O*(1), whereas the production and decay rates are *O*(1*/ε*), where *ε* ≪ 1 is a small dimensionless parameter. The aim of the paper is to find asymptotic approximations, valid for *ε* ≪ 1, of the stationary behaviour of the model.

Let us talk through the specific forms of the reaction rates and reset maps of the individual reactions in Table 1. The production rate depends on the number *s* of active protein through a general (integer-valued) feedback response function *f*_*s*_. The production reset map indicates that the number *X* of inactive protein is increased by the size *B* ≥ 1 of a production burst. Bursts sizes are drawn (independently of each other) from a distribution

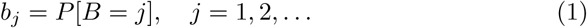

The activation rate is proportional to the number of inactive protein; the factor of proportionality is set to one without loss of generality. The activation reset map turns one inactive protein into an active protein if there is extra capacity for active protein (*s* < *s*_max_); it removes an inactive protein without creating an active protein if there is no capacity (*s* = *s*_max_). The importance of introducing an upper bound *s*_max_ on the number of active protein is discussed at greater length in Section 7. The decay rate is proportional to the number of active protein; the decay reset map decreases the number of active protein by one.

## 3 QSS approximation

In the slow-activation regime (*ε* ≪ 1), the inactive protein X is present at a copy number *X* (written in italics) which is *O*(*ε*^−1^) large. In order to measure the abundance of species X on an *O*(1) scale, we define

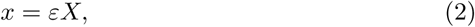

which we refer to as the concentration of the inactive protein X. In the limit of *ε* → 0, the concentration *x* becomes a continuous quantity.

An additional effect of small values of *ε* is that the turnover of the active protein S becomes much faster than that of the inactive protein X. This difference in timescales leads to seeking a quasi-steady-state (QSS) reduction of the model. On the fast timescale of the turnover of S, we treat the concentration *x* of the slow species X as a fixed quantity. With this assumption, the dynamics of S is that of an *M/M/s*_max_*/s*_max_ server with memoryless arrival and service times, *s*_max_ servers, and no queue (Gross, 2008). This analogy identifies active protein molecules S with customers, protein activation with customer arrival (occurring with a constant rate *ε*^−1^*x*), and protein decay with customer service (with rate *ε*^−1^ per customer). A well-known result from the queuing theory implies that the QSS distribution of *s* is

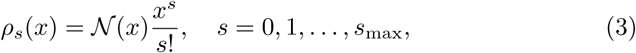

where

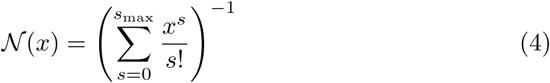

is the normalisation constant. Thus, fixing the slow species X to a concentration *x*, the fast species copy number *s* obeys the (truncated) Poisson distribution (3) with location parameter *x*.

Given a concentration *x* of the inactive protein X, the effective production burst rate is equal to 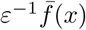, where 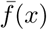 is the expectation of *f*_*s*_ with respect to the QSS distribution (3), i.e.

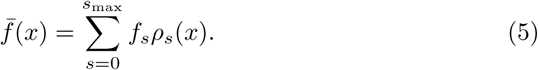

The value of 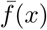 is determined by all values of *f*_*s*_: for example, 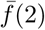 includes, with a small weight, the value *f*_10_; conversely, *f*_2_ contributes a little towards 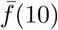. Nevertheless, the main contribution to 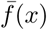 comes from the values of *f*_*s*_ whose argument *s* is close to *x*. Due to the contributions of neighbouring terms, any sharp features of *f*_*s*_ are “mollified” in the function 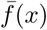; for example, a step function (Gedeon et al, 2017; Crawford-Kahrl et al, 2019)

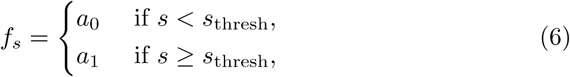

tunrs into a smooth sigmoid function by the application of (5); see Figure 1, top panels.

**Figure 1:**
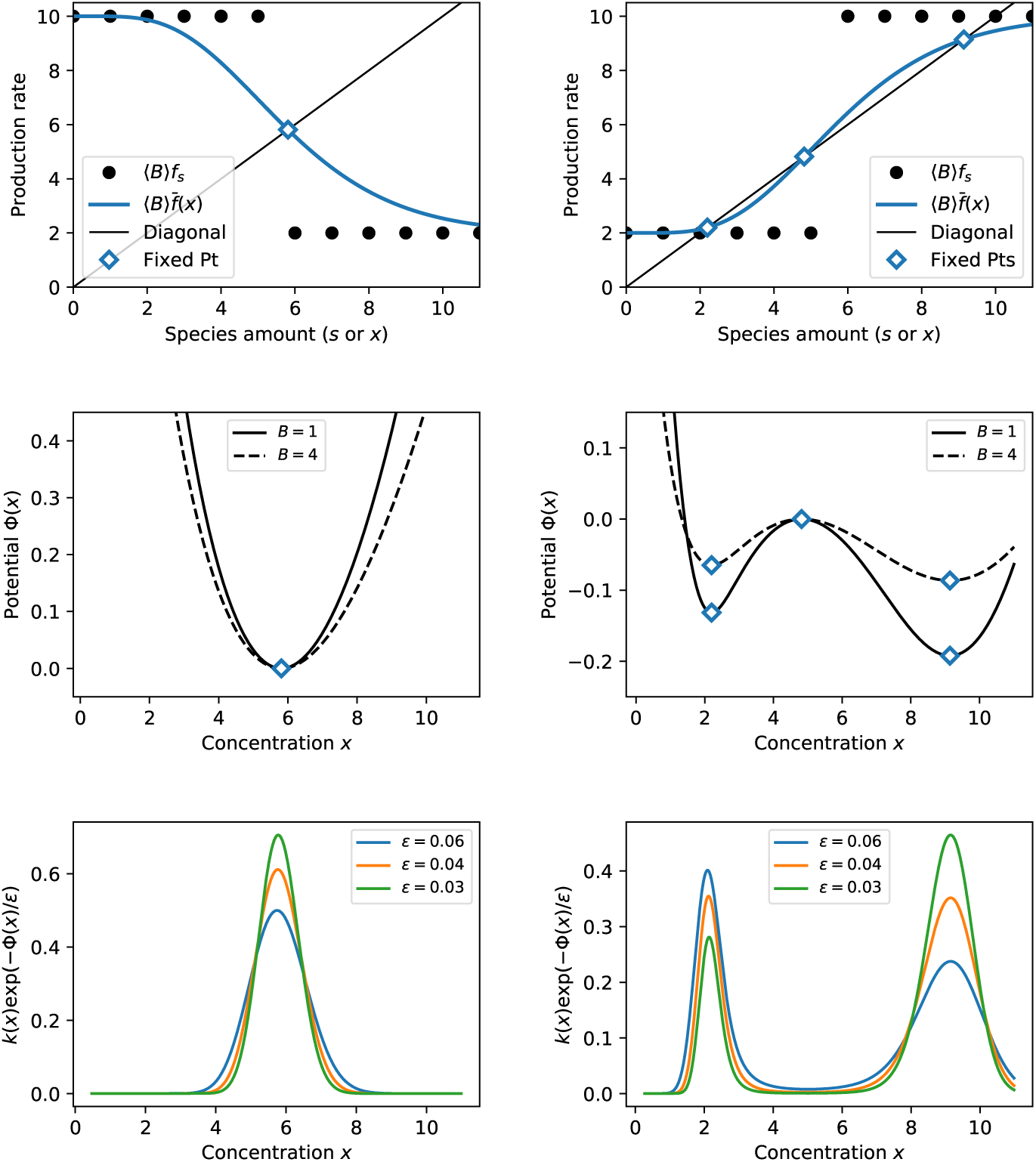
*Top:* The instantaneous production rate ⟨*B*⟩ *f*_*s*_ (black circles) and the QSS-averaged production rate 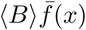 (blue line) for negative (*left*) or positive (*right*) feedback. Blue diamond markers indicate the fixed points. *Middle:* The WKB potential *Φ*(*x*). Its local extrema colocate with the fixed points of the QSS-averaged production rate. The potential gets flatter as production becomes bursty (dashed line). *Bottom:* The WKB approximation of the probability density function of the inactive protein concentration *x*. The pdf is peaked around the potential minima for *ε* ≪ 1. *Parametric values:* The upper bound on *S* is *s*_max_ = 20. The feedback threshold is *s*_thresh_ = 6. The burst size is fixed to *B* = 1 except for the dashed line, middle panels, where it is fixed to *B* = 4. Unimodal examples use *a*_0_ = 10, *a*_1_ = 2, except for the dashed line, middle panels, which uses *a*_0_ = 2.5, *a*_1_ = 0.5. Bimodal examples use *a*_0_ = 2, *a*_1_ = 10, except for the dashed line, middle panels, which uses *a*_0_ = 0.5, *a*_1_ = 2.5.

The concentration of protein produced per unit time is equal to the product of the effective burst rate 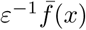 and the mean burst size *ε* ⟨*B*⟩ (in units of concentration), where

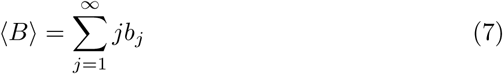

is the mean burst size (in units of copy number). Subtracting from the product 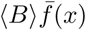 the linear activation rate yields the limiting deterministic dynamics

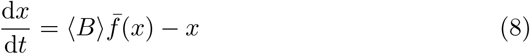

for the inactive protein. Steady states of (8) are the intersections of 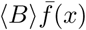 with the diagonal (i.e. fixed points), and we consider two cases separately:

### Monostable case

Assume that (8) possesses a single globally stable steady state *x*_0_ (Figure 1, top left, blue diamond). After the elapse of an initial transient, the inactive protein concentration *x* is attracted to *x*_0_, and the active protein copy number *s* follows the QSS distribution *ρ*_*s*_(*x*_0_).

### Bistable case

Assume that (8) possesses three steady states *x*_−_ < *x*_0_ < *x*_+_, of which *x*_−_ and *x*_+_ are stable and *x*_0_ is unstable (Figure 1, top right, blue diamonds). Since in the long run the inactive protein concentration will alternate between being very close to either of the two attractors, the steady-state distribution of the active protein is expected to take the form of a mixture of the QSS distributions *ω*_−_*ρ*_*s*_(*x*_−_) + *ω*_+_*ρ*_*s*_(*x*_+_), where *ω*_−_ ≥ 0, *ω*_+_ ≥ 0, *ω*_−_ + *ω*_+_ = 1, are the probabilistic weights of the stable steady states *x*_−_ and *x*_+_. Thus, we expect the steady-state distributions of *s* to be a mixture of two Poissonian modes located at the two stable steady states of the ODE (8).

## 4 WKB approximation: an overview

The approximation introduced in this Section reinforces the QSS-based heuristics with a systematic argumentation. Additionally, it provides asymptotic expansions as *ε* → 0 of the weights in the mixture distributions (which the QSS heuristics does not).

The WKB approximation of the probability mass function *p*(*x, s*; *ε*) of the inactive protein concentration *x* and the active protein copy number *s* is sought (and found in Section 6) in the form of

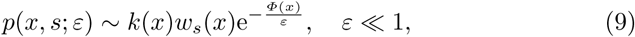

where *k*(*x*), *w*_*s*_(*x*) and *Φ*(*x*) are independent of *ε* and satisfy *k*(*x*) > 0, *w*_*s*_(*x*) > 0, and 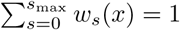. The variable *s* is placed in the subscript to emphasise its discreteness (contrasting it with the continuous nature of *x* in the limit of *ε* → 0). The ansatz (9) can be rephrased in terms of the marginal distribution of *x* and the conditional distribution of *s* as

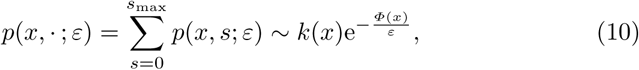

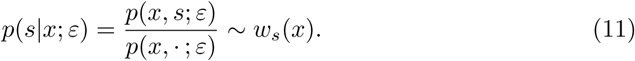

We refer to *Φ*(*x*) as the WKB potential, to *k*(*x*) as the WKB prefactor, and to *w*_*s*_(*x*) as the WKB conditional distribution of *s*. In Section 6 we determine these terms from the underlying chemical master equation and establish a connection between the WKB and QSS approximations. At this stage, let us highlight the main outcomes of the forthcoming analysis.

We will show that the potential is a Lyapunov function of the limiting ODE model (8), possessing local minima (maxima) at the same points where (8) has stable (unstable) steady states (Figure 1, centre). Additionally, the potential carries information about the noise in the model that the ODE does not: specifically, the ODE depends only on the product of burst rate and burst frequency, remaining the same if the burst size is multiplied by the same factor as the burst frequency is divided by; the potential, on the other hand, becomes flatter as the system becomes more bursty (Figure 1, centre, dashed lines). This observation is consistent with an intuition that bursty production enhances noise and the chance to escape potential wells.

If the ODE is monostable, the potential has a single global minimum situated at the single global attractor *x*_0_ of the differential equation (Figure 1, centre left). For *ε* ≪ 1 the distribution (10) of *x* is sharply peaked around *x*_0_ (Figure 1, bottom left) so that it can be approximated by a delta peak

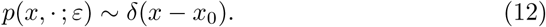

If the ODE is bistable, *Φ*(*x*) is a double-well potential, with the wells at the stable steady states *x*_−_ and *x*_+_ being separated by a barrier at the unstable steady state *x*_0_ (Figure 1, centre right). For *ε* ≪ 1 the distribution (10) of *x* is sharply peaked around *x*_−_, *x*_+_ (Figure 1, bottom right) so that a mixture of delta peaks

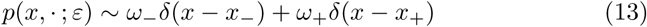

can be used as a rough approximation.

We will show in Section 6 that at a steady state *x*_*_ ∈ {*x*_−_, *x*_0_, *x*_+_} of (8) the WKB conditional distribution of *s* coincides with the QSS approximation,

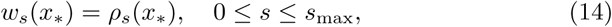

where *ρ*_*s*_(*x*) is defined by (3)–(4). Interestingly, conditioning on points *x* away from the steady states of (8) leads to WKB conditional distributions of *s* that differ from the QSS heuristics. Arguably, this disagreement arises because the simplifying QSS assumption of a fixed *x* is invalidated by making a large deviation from a steady state. Nevertheless, the contribution of non-QSS conditional distributions towards the total distribution of *s* turns out to be negligible.

The total distribution of *s* is given by

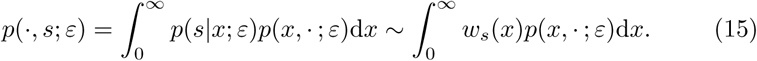

Inserting the delta-peak singleton (12) or the delta-peak-mixture (13) approximations into (15) and using (14) we find a Poisson singleton

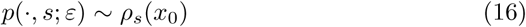

for the monostable case and a Poisson mixture

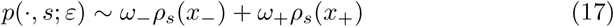

for the bistable case. These are the very approximations which were suggested by the QSS approach.

For the weights *ω*_−_ and *ω*_+_ we have

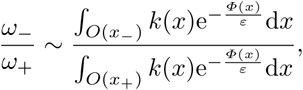

where *O*(*x*_−_) and *O*(*x*_+_) are neighbourhoods of *x*_−_ and *x*_+_. Applying the Laplace method to approximate for small *ε* the integrals in the above equation, we find

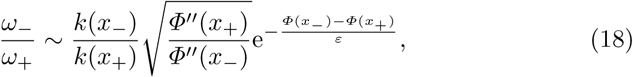

which together with *ω*_−_ + *ω*_+_ = 1 determine the asymptotic behaviour of the weights. In particular, if the right well of the potential *Φ*(*x*) is deeper than the right well, i.e. *Φ*(*x*_−_) > *Φ*(*x*_+_), then (18) implies that (*ω*_−_, *ω*_+_) → (0, 1) as *ε* →0 (and (*ω*_−_, *ω*_+_) → (1, 0) as *ε* → 0 if *Φ*(*x*_−_) < *Φ*(*x*_+_)). However, if the two wells are finely balanced, the weights are comparable for a range of *ε* ≪ 1, and both Poissons that constitute the steady-state distribution (17) of *s* are appreciable.

The approximation of the distribution of *x* by a single delta peak (monostable case) or a mixture of two delta peaks (bistable case) is sufficient for the purpose of deriving the leading-order approximation of the distribution of *s*. If one is interested in the distribution of *x* itself, a satisfactory result is obtained by a parabolic approximation of the WKB potential around its minima. In the monostable case it is given by a Gaussian singleton

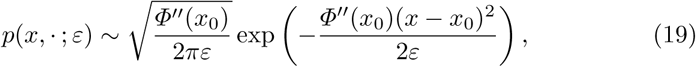

and in the bistable case by a Gaussian mixture

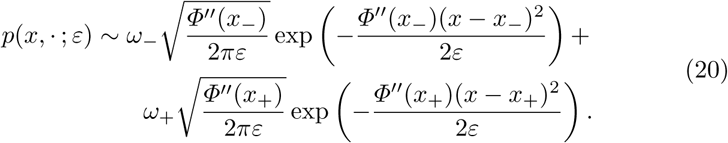

The results (19) and (20) for the inactive protein distribution are expressed in terms of the concentration *x*. Expressions in terms of the copy number *X* = *x/ε* can be obtained using the well-known rule for a linear transformation of a normally distributed random variable.

For further background on this Section’s results as well as practical recipes for the calculation of the WKB terms that appear in these results, we refer the reader to Section 6. In the next Section, we introduce the master equation, find its steady-state solution numerically, and compare it to the Poisson/Gaussian singleton/mixture approximations.

## 5 Master equation

The probability *P* = *P*(*X, s, t*) of having *X* molecules of inactive protein X and *s* molecules of active protein S (cf. Table 1) at time *t* satisfies the chemical master equation (CME)

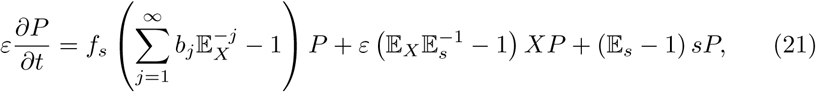

in which 𝔼_*X*_ and 𝔼_*s*_ denote the van Kampen step operators (van Kampen, 2006) in variables *X* and *s* and 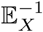 and 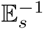 are their formal inverses. The master equation (21) applies in the unbounded case *s*_max_ = ∞; the bounded case will be dealt with in detail in Section 6. The principal interest of this paper is in steady-state solutions, which are obtained by setting the derivative in (21) to zero.

The first step in finding a numerical solution to a CME is its truncation to a finite number of equations. Following the approach illustrated e.g. by Borri et al (2016), we truncate the master equation to a finite lattice {0, 1, …, *s*_max_}×{0, 1, …, *X*_max_}, and calculate the (unique) normalised steady-state solution; this amounts to finding a nullvector of a sparse square matrix of large order (*s*_max_ +1)(*X*_max_ +1). The upper bound for the active protein is set to *s*_max_ = 20, while the upper bound *X*_max_ for the inactive protein is set to *X*_max_ = 4 [*x*_+_*/ε*], where *x*_+_ is the uppermost steady state of the ODE (8). Preliminary rounds of the Gillespie stochastic simulation algorithm (SSA) (Gillespie, 1976) show that sample trials (almost) never exceed *X*_max_, so that the truncated solution is expected to be a good approximation of the original one.

The numerical and the WKB-based asymptotic solutions to the CME are in a good agreement both for negative feedback (Figures 2–3) as well as for positive feedback (Figure 4–5). For each feedback type we consider two (fixed) burst sizes *B* = 1 and *B* = 4 (*b*_*B*_ = 1, *b*_*j*_ = 0 for *j* ≠ *B*). Panel columns in the figures refer to the two protein species; panel rows refer to the chosen values of *ε*. The parameters of the step function (6) are given in the figure captions. The asymptotic approximations (Figures 2–5, blue lines) are obtained by evaluating (16) and (19) in the monostable case (negative feedback) or (17) and (20) in the bistable case (positive feedback). The Poisson modes are evaluated from the formula (3)–(4). The weights in the mixture distributions are calculated from (18). The numerical values of the coefficients that enter these calculations are summarised in Table 2.

**Table 2:** Coefficients of the WKB approximation used in the numerical examples of Figures 2–5. The values of *x*_*_ give the location of the stable fixed points of ODE (8). The value of (*Φ*″(*x*_*_))^−1^ gives (up to a factor of 1*/ε*) the variance of the Gaussian modes. The prefactor ratio *k*(*x*_−_)*/k*(*x*_+_) and the potential difference *Φ*(*x*_−_) − *Φ*(*x*_+_) enter into the calculation of the mixture weights in the bistable case.

| Burst size | Feedback | $x_*$ | $(\Phi''(x_*))^{-1}$ | $k(x_-)/k(x_+)$ | $\Phi(x_-) - \Phi(x_+)$ |
| --- | --- | --- | --- | --- | --- |
| 1 | Negative | 5.81 | 10.69 | N/A | N/A |
| 4 | Negative | 5.81 | 14.44 | N/A | N/A |
| 1 | Positive | 2.2 | 3.35 | 4.47 | 0.0606 |
|  |  | 9.14 | 14.98 |  |  |
| 4 | Positive | 2.2 | 8.67 | 4.42 | 0.0216 |
|  |  | 9.14 | 40.17 |  |  |

**Figure 2:**
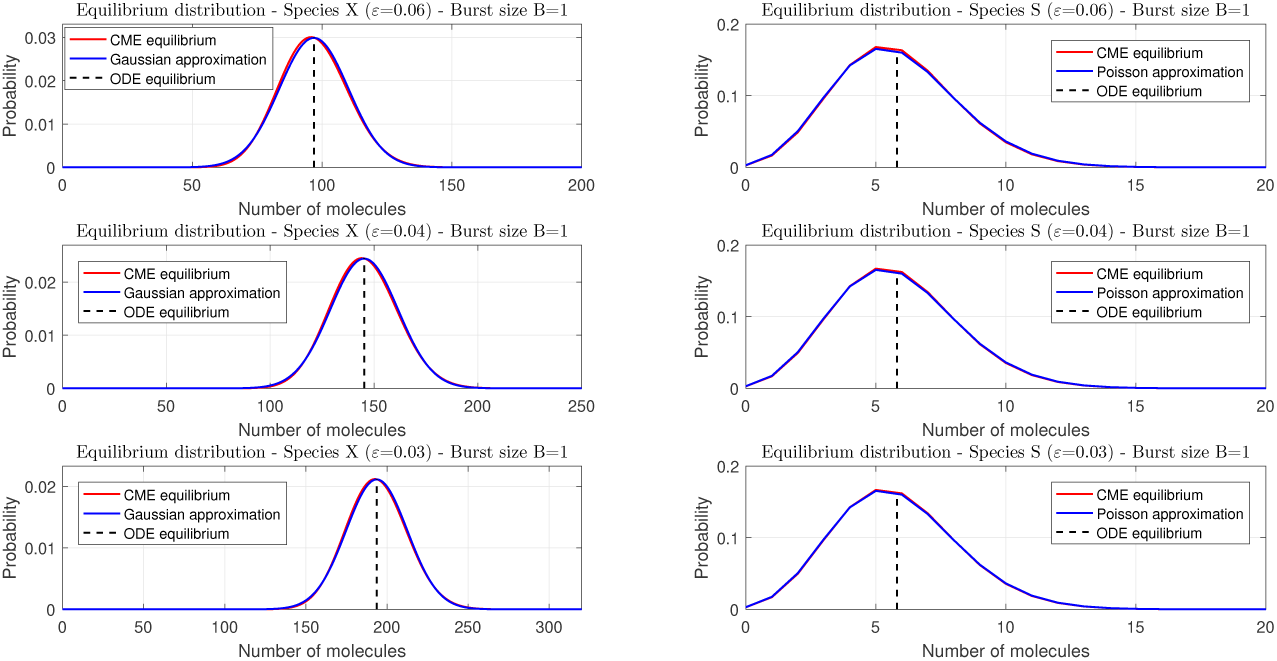
Steady-state distributions for negative feedback and fixed burst size *B* = 1 obtained by numerical solution (red line) and asymptotic approximation (blue line). Panel columns refer to the two protein species; panel rows refer to distinct values of *ε*. The dashed lines indicate the locations of stable fixed points of the deterministic rate equation. The step function (6) parameters are *s*_thresh_ = 6, *a*_0_ = 10, *a*_1_ = 2.

**Figure 3:**
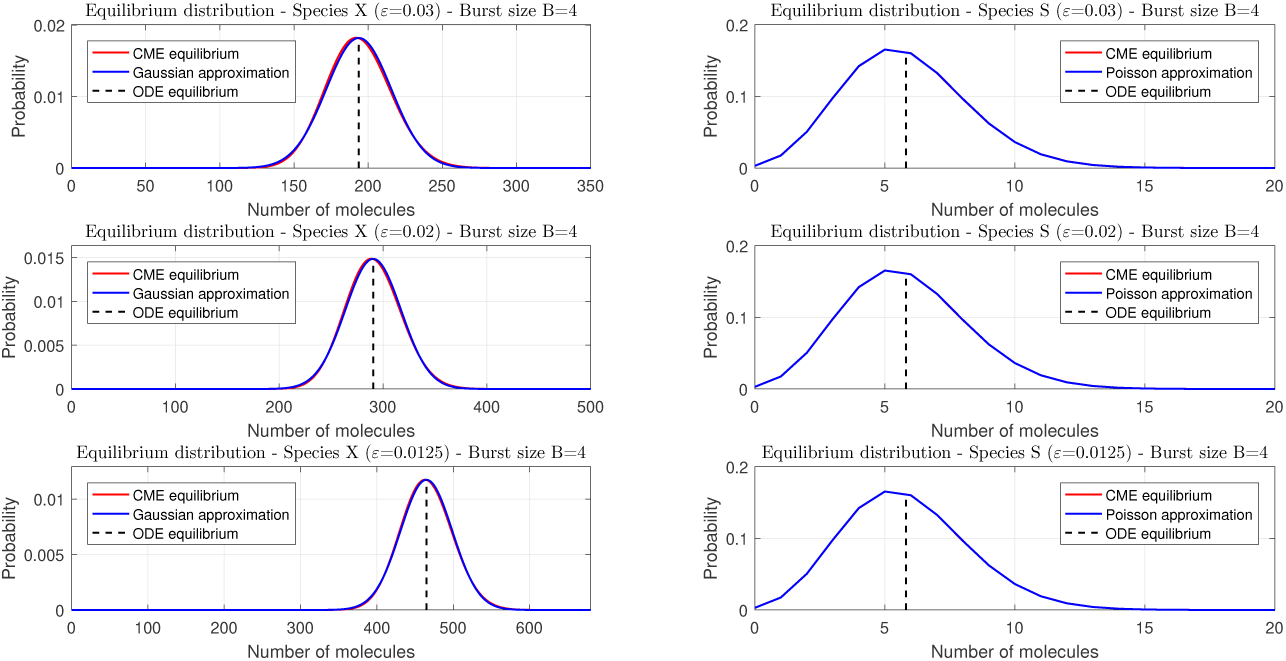
Steady-state distributions for negative feedback and fixed burst size *B* = 4 obtained by numerical solution (red line) and asymptotic approximation (blue line). Panel columns refer to the two protein species; panel rows refer to distinct values of *ε*. The dashed lines indicate the locations of the stable fixed points of the deterministic equation. The step function (6) parameters are *s*_thresh_ = 6, *a*_0_ = 2.5, *a*_1_ = 0.5.

**Figure 4:**
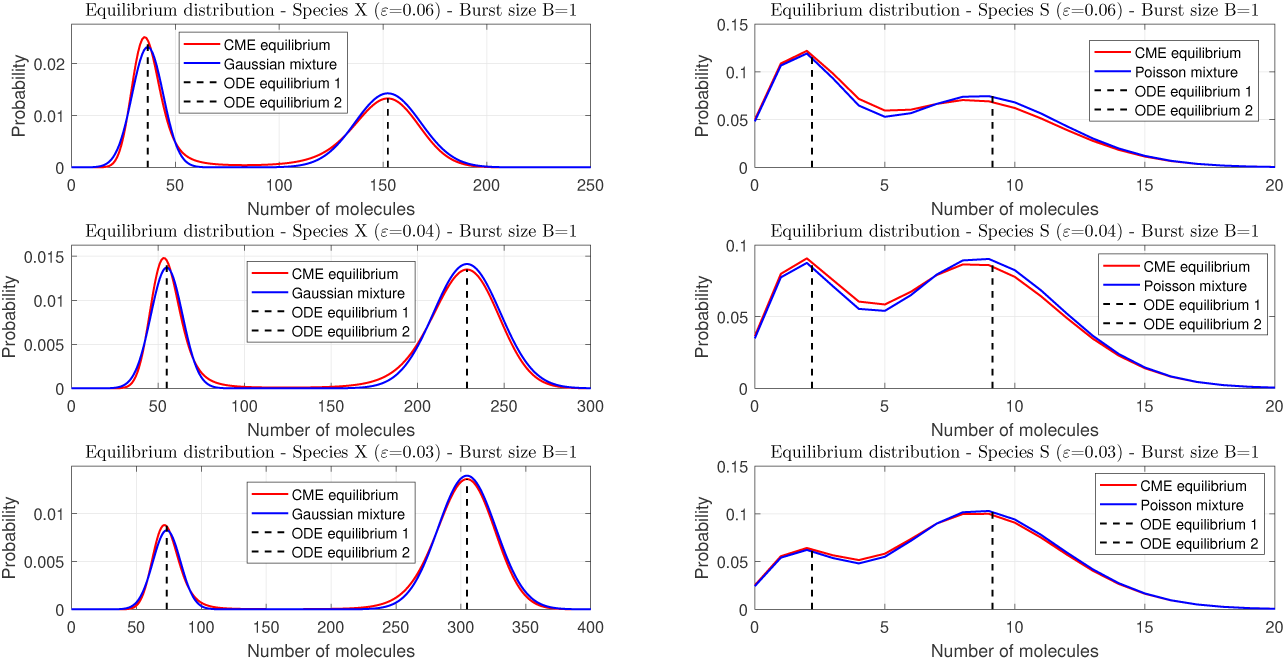
Steady-state distributions for the case of positive feedback and fixed burst size *B* = 1 obtained by numerical solution (red line) and asymptotic approximation (blue line). Panel columns refer to the two protein species; panel rows refer to distinct values of *ε*. The dashed lines indicate the locations of the stable fixed points of the deterministic equation. The step function (6) parameters are *s*_thresh_ = 6, *a*_0_ = 2, *a*_1_ = 10.

**Figure 5:**
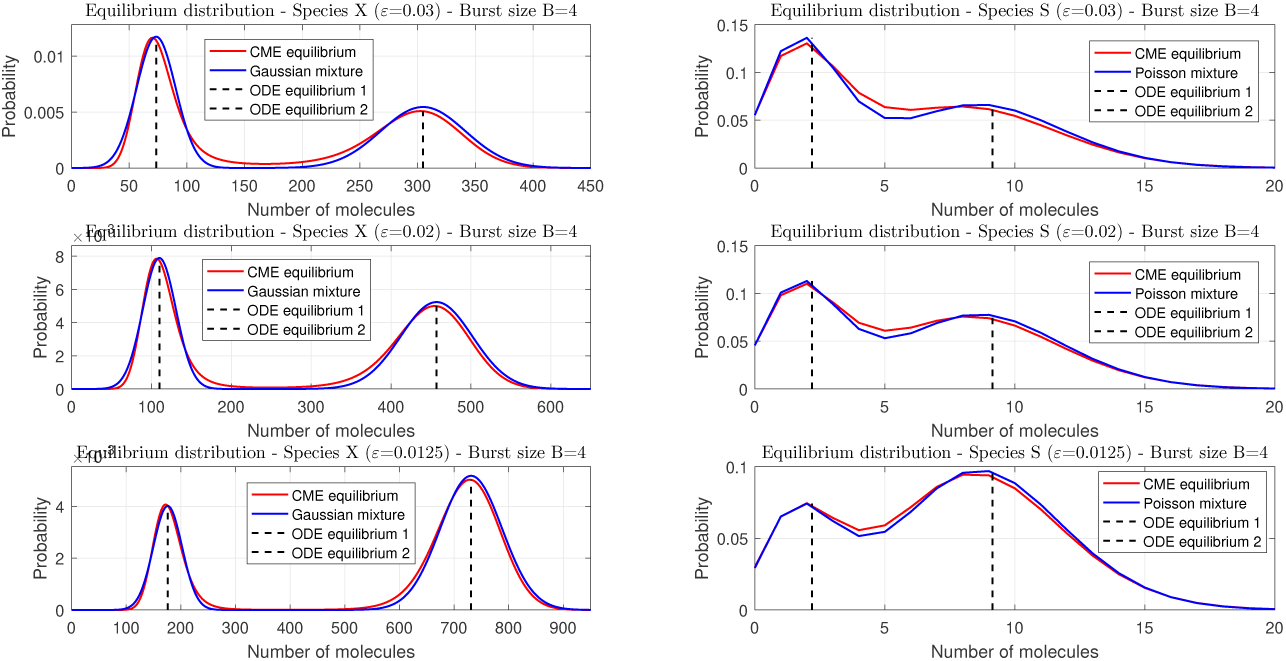
Steady-state distributions for the case of positive feedback and fixed burst size *B* = 4 obtained by numerical solution (red line) and asymptotic approximation (blue line). Panel columns refer to the two protein species; panel rows refer to distinct values of *ε*. The dashed lines indicate the locations of the stable fixed points of the deterministic equation. The step function (6) parameters are *s*_thresh_ = 6, *a*_0_ = 0.5, *a*_1_ = 2.5.

## 6 WKB approximation: analysis

We start our analysis by formulating a version of the master equation (21) that is applicable to the *s*_max_ < ∞ case (cf. Table 1). For this purpose it turns out to be helpful to use a modified version of the van Kampen step operators. For a sequence *v*_*s*_ indexed by an integer 0 ≤ *s* ≤ *s*_max_, we define the left- and right-shift operators by

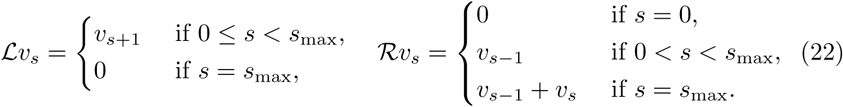

Away from the boundary *s* = *s*_max_, the left- and right-shift operators ℒ and ℛ are equal to the van Kampen step operator and its formal inverse, respectively. Let us discuss the meaning of the modifications of the operators that are made on the boundary *s* = *s*_max_. The left-shift operator is used below to describe the transfer of probability mass due to protein decay. The modification of the left-shift operator at the boundary *s* = *s*_max_ means there is no transfer of probability from the inadmissible state *s*_max_ + 1. The right-shift operator is used below to describe the transfer of probability mass due to protein activation. The reset map of the activation reaction channel (Table 1) implies that the states with *s*_max_ as well as the states with *s*_max_ - 1 active protein molecules transfer into a state with *s*_max_ molecules. Correspondingly, the right-shift operator returns at the boundary the sum of the ultimate and the penultimate terms of the original sequence.

The probability *P*(*X, s, t*) of at time *t* having *X* molecules of inactive protein X and *s* molecules of active protein S (cf. Table 1) satisfies

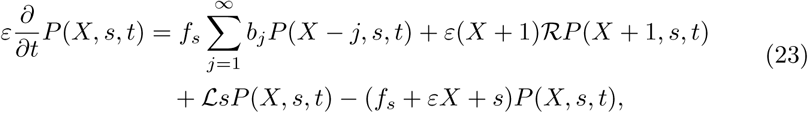

in which we use the operator formalism (22) in the variable *s* but the shifts in the variable *X* are made explicit. We thereby tacitly understand, as is customary in analyses of chemical master equation, that the probability of having a negative number of species *X* is equal to zero.

Inserting

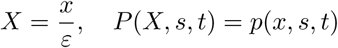

into (23), we obtain

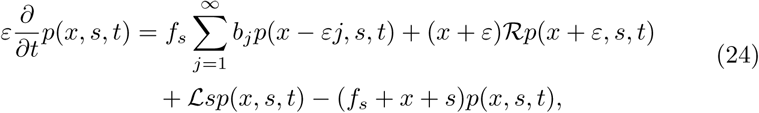

which expresses the master equation in terms of the concentration of the inactive protein. Equating the derivative in (24) to zero, we arrive at a bivariate difference equation

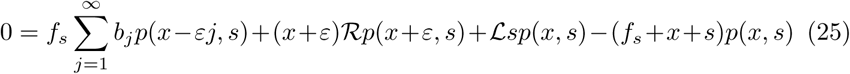

for the steady-state distribution *p*(*x, s*). Below we seek an asymptotic approximation as *ε* → 0 to the (normalised) solution of (25).

### 6.1 Expansion

We seek a solution to (25) in the WKB form, cf. Equation (9),

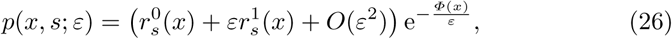

where 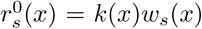 and 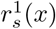 give the first two terms in the asymptotic expansion in the powers of *ε*. Below we develop, by means of (26), the individual terms of the difference equation (25) into asymptotic expansions of up to the second order. This is a mechanistic but laborious exercise. Therefore we suggest that, on first reading, the reader focus their attention on the leading-order terms in the expansions; these are sufficient for evaluating the potential *Φ*(*x*) and *w*_*s*_(*x*) (which play the central role in the analysis). The second-order terms, including those that involve 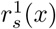, come into play in SubSection 6.3 in the determination of the prefactor *k*(*x*).

For the first term in equation (25) we find

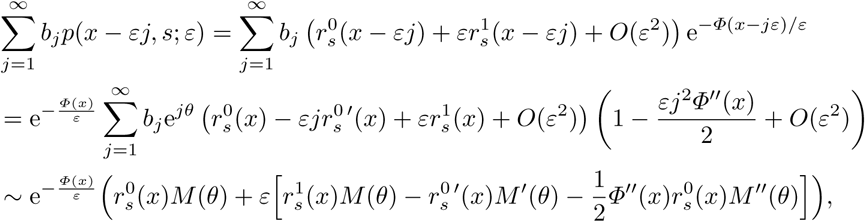

where

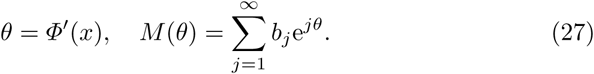

are the potential derivative and the moment generating function of the burst-size probability distribution (1), respectively.

The second term in equation (25) is developed into

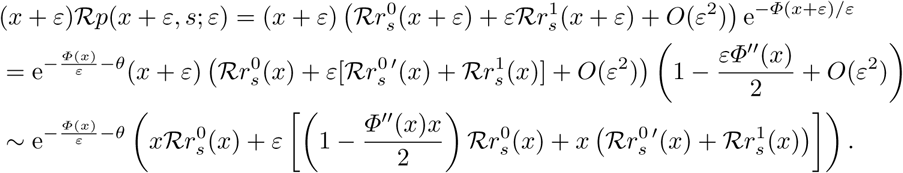

The remaining terms in (25) are easy to expand.

We insert the WKB ansatz (26) into (25), expand the individual terms of the equation as suggested above, and collect terms of same order; at the leading order this yields

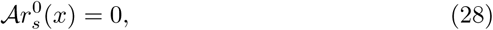

where

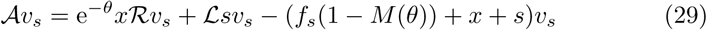

is a linear operator acting on sequences *v*_*s*_ defined for 0 ≤ *s* ≤ *s*_max_. Such sequences can be represented by (*s*_max_ + 1)-dimensional column vectors 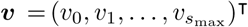, and the linear operator 𝒜 as a square matrix ***A*** of order *s*_max_ + 1. Here and throughout this Section, we will go back and forth between the operator–sequence and the matrix–vector notations, using that which expresses a given formula more succinctly.

The matrix ***A*** is tridiagonal. On the diagonal it has the sequence −*f*_*s*_(1 − *M* (*θ*)) − *x* − *s*, where 0 ≤ *s* < *s*_max_, except for the last diagonal element, which is given by 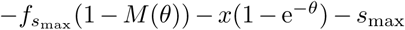. On the upper diagonal it has the sequence 1, 2, …, *s*_max_; the elements of the lower diagonal are all equal to e^−*θ*^*x*. It looks like

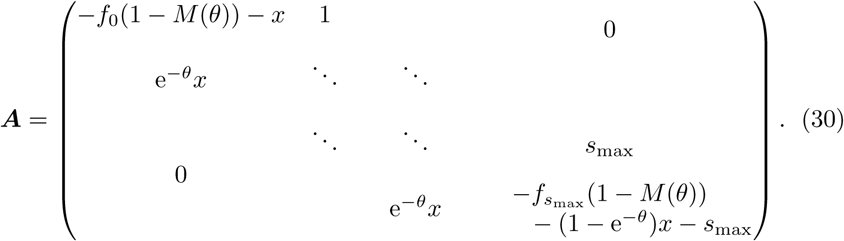

The matrix ***A*** = ***A***(*x, θ*) — just like the associated operator 𝒜= 𝒜 (*x, θ*) — depends on the protein concentration *x* and the (yet unknown) potential derivative *θ* = *Φ*′(*x*).

Equation (28) can be written in a matrix form as

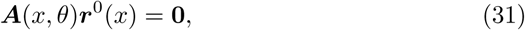

where 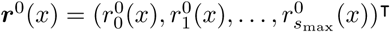 is a positive vector (i.e. a vector with only positive elements). The matrix ***A*** does not have any negative off-diagonal elements. Denote by

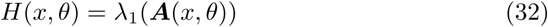

the eigenvalue with largest real part (which we refer to hereafter as the principal eigenvalue). The Perron–Frobenius theorem implies that the principal eigenvalue is real and that its right and left eigenvectors (referred to as principal eigenvectors) are real and positive. The right eigenvectors of non-principal eigenvalues, being orthogonal to the left principal eigenvector, cannot be positive. Therefore, equation (31) is solvable in ***r***^0^(*x*) > 0 if

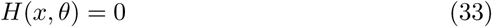

holds. Solving equation (33) in *θ*, we obtain a functional dependence of the potential derivative *θ* = *Φ*′(*x*) on the concentration *x*. The potential itself can be obtained by (numerically) integrating its derivative.

Crucially, equation (33) does not define *θ* = *θ*(*x*) uniquely. First, there is a trivial solution *θ* = 0 for any value *x* > 0. Indeed, *θ* = 0 implies *M* (0) = 1, with which the matrix ***A***(*x*, 0) reduces to (the transpose of) the transition rate matrix of the QSS process (the *M/M/s*_max_*/s*_max_ server; see Section 3). Therefore it is singular and its principal eigenvalue (32) is equal to zero. The trivial branch *θ* ≡ 0 of solutions to (33) cannot be used to construct the potential: we need to look for a different branch. To do so, it is useful to consider a Hamiltonian system of differential equations corresponding to the Hamiltonian (32).

### 6.2 The Hamiltonian system

Differentiating with respect to *x* equation (33) in which *θ* = *θ*(*x*) yields

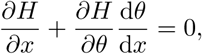

i.e.

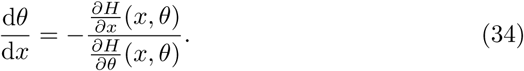

The non-autonomous differential equation (34) is equivalent to the system of two autonomous differential equations

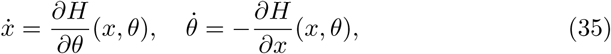

in which the dot represents the time derivative. The system is Hamiltonian: its trajectories form the level sets of (32). We are specifically interested in the zero set (33). Borrowing terminology from the Hamiltonian formalism, we refer to the variable *θ* as the conjugate momentum.

In order to solve system (35), we need to evaluate the right-hand sides. For this purpose it is useful to express the Hamiltonian (32) as

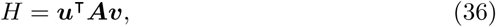

where ***u*** > 0 and ***v*** > 0 are the left and right eigenvectors corresponding to the principal eigenvalue *H*, i.e.

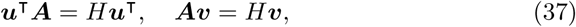

which additionally satisfy

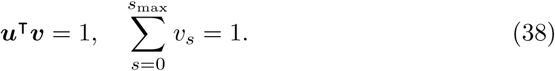

Conditions (38) can be met by a suitable choice of multiplicative constants.

The principal eigenvectors ***u*** and ***v***, just like ***A*** and the principal eigenvalue *H*, depend on the protein concentration *x* and the conjugate momentum *θ*. For *θ* = 0 the matrix ***A***(*x*, 0) reduces to the transition rate matrix of the QSS process with principal eigenvalue *H*(*x*, 0) = 0; the principal eigenvectors are given by the nullvectors of the QSS process

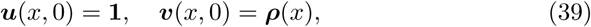

in which **1** = (1, 1, …, 1)^**T**^ is an (*s*_max_ + 1)-dimensional vector of ones and 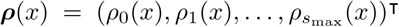 is the vector representation of the QSS stationary distribution (3).

The partial derivatives of the Hamiltonian (36) are given by

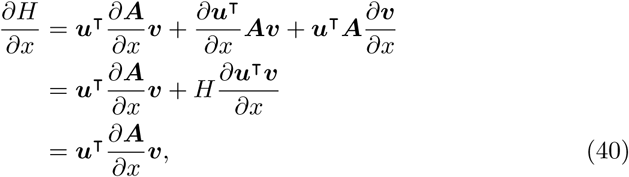

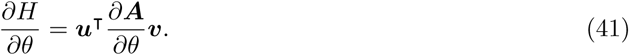

The derivatives of ***A*** in (40)–(41) are matrix representations of the derivatives

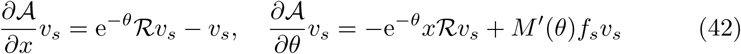

of the operator 𝒜 (29).

For *θ* = 0 the derivatives (40)–(41) of the Hamiltonian satisfy

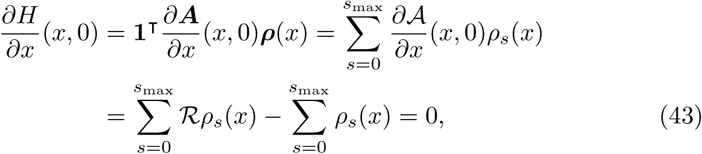

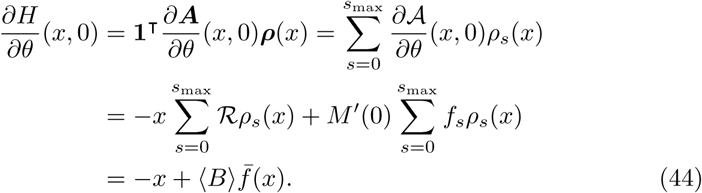

We note that in the equalities leading up to (43) and (44), we made use of the following properties: (i) the product of a row vector of ones and a column vector is equal to the sum of the column vector’s elements; (ii) the right-shift operator ℛ (22) preserves the sum of vector elements; (iii) evaluating at zero the first derivative of the burst-size moment generating function (27) gives the mean burst size; we also used the definition of the QSS-averaged production rate (5).

Inserting *θ* = 0 into (35) and using (43)–(44), we find that on the *x*-axis the Hamiltonian system satisfies

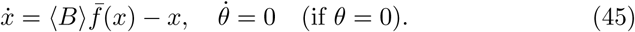

Equations (45) imply that (i) the *x*-axis is an invariant set of the Hamiltonian system and that (ii) the system reduces to the deterministic rate equation (8) when restricted to the invariant set. The Hamiltonian system (35) thus comprises, and extends by an additional dimension in *θ*, the ODE dynamics established by the quasi-steady-state analysis of Section 3.

A point (*x*_*_, 0), where *x*_*_ ∈ {*x*_−_, *x*_0_, *x*_+_} is any of the fixed points of (8), is also a steady state of the full Hamiltonian system (35). The linearisation matrix is given by

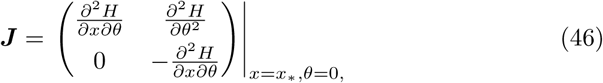

in which the second derivative of *H* with respect to *x* is immediately seen to be zero because of (43).

It follows that (*x*_*_, 0) is a saddle of (35) with eigenvalues

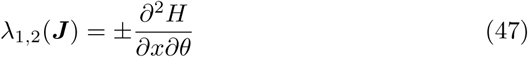

and eigenvectors

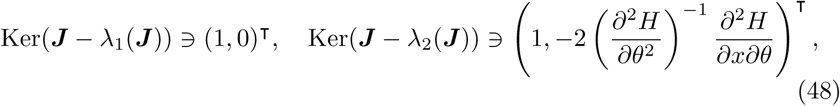

in which the derivatives of the Hamiltonian are evaluated at (*x*_*_, 0). One can show that

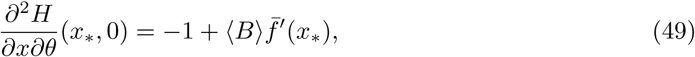

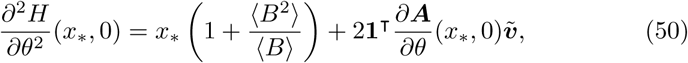

where 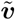 is a solution to 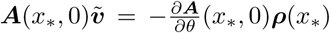. Equation (49) is an immediate consequence of (44). Equation (50) is derived in Appendix A. The derivative of the effective production rate (5) satisfies

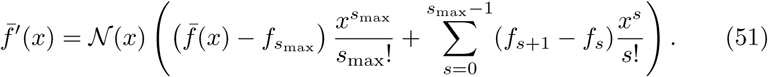

Equations (49)–(51) provide a practical numerical recipe for calculating the nontrivial eigenvector (48) of the Hamiltonian system linearisation.

The trajectories emanating from a saddle (*x*_*_, 0) along the direction of the eigenvector (1, 0)^**T**^ form the trivial branch *θ* = 0 of the zero set (33). The trajectories emanating from the saddle along the nontrivial eigenvector (48) form the nontrivial branch of the zero set (33) (Figure 6, left). The nontrivial branch constitutes the sought-after WKB potential derivative *θ* = *Φ*′(*x*). The derivative *Φ*′(*x*) is equal to zero where the nontrivial *θ* = *Φ*′(*x*) and the trivial *θ* = 0 branches intersect, i.e. at the fixed points *x* = *x*_*_ ∈ {*x*_−_, *x*_0_, *x*_+_}of the deterministic rate equation (8). This establishes the claim made in Section 4 that the potential extrema coincide with the fixed points of the deterministic model.

**Figure 6:**
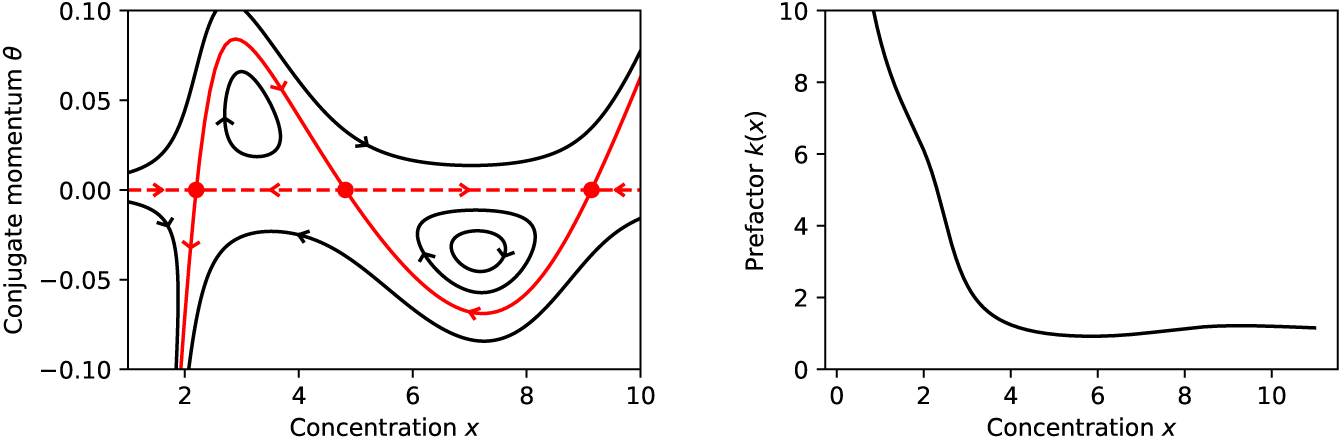
*Left:* Phase plane of the Hamiltonian system (35). The heteroclinic orbits (red colour) form the zero set of the Hamiltonian (32). The nontrivial portion of the zero set (full red line) defines the derivative of the WKB potential *θ* = *Φ*′(*x*). The trivial portion of the zero set (*θ* = 0; dashed red line) is the phase line of the deterministic ODE (8). The red markers denote the steady states of (8), which are also saddles of the Hamiltonian system. *Right:* The WKB prefactor. The parameter values are: The upper bound on *S* is *s*_max_ = 20. The burst size is fixed to *B* = 1. The step function (6) parameters are *s*_thresh_ = 6, *a*_0_ = 2, *a*_1_ = 10.

Formulae (18), (19), and (20) require us to evaluate the second derivative of the potential at fixed points. The second derivative of *Φ*(*x*), i.e. the first derivative of *θ* = *Φ*′(*x*), is equal at the fixed points to the second component of the nontrivial eigenvector (48) of the Hamiltonian system linearisation (cf. Figure 6, left panel). Away from fixed points, the derivative of *θ* = *Φ*′(*x*) can be evaluated by substituting into the right-hand side of (34).

In order to complete the WKB approximation (9) at the leading order, we express

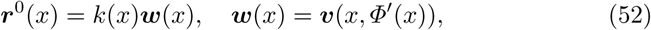

in which the *l*^1^-normalised nullvector 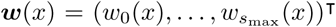 gives the distribution of *s* conditioned on a concentration *x*, and *k*(*x*) is a prefactor. In particular, if *x* = *x*_*_ is a fixed point, then we have ***w***(*x*_*_) = ***v***(*x*_*_, 0) = ***ρ***(*x*_*_) (the Poissonian QSS distribution); this establishes the claim (14) made in Section 4. Away from the fixed points, the conditional probability mass function of *s* is in general different from the Poissonian QSS distribution.

Having determined the potential *Φ*(*x*) and the probability mass function *w*_*s*_(*x*), the last remaining task is to calculate the prefactor *k*(*x*). Doing so requires us to consult the second-order terms in the WKB expansion.

### 6.3 Calculating the prefactor

Having inserted the WKB ansatz (26) into the master equation (25) and having expanded the individual members up to the second order (Section 6.1), we now collect the second order terms and obtain

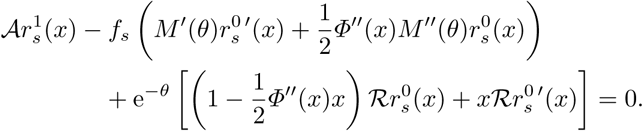

Inserting the factorisation 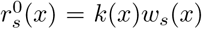, cf. (52), into the equation above yields

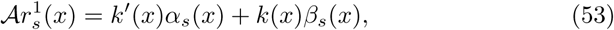

where

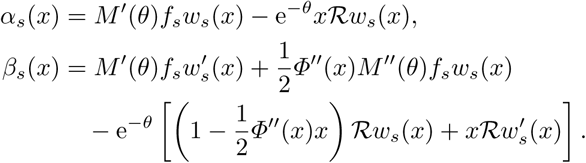

In order that equation (53) be solvable in 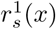, its right-hand side must be orthogonal to the left nullvector *l*_*s*_(*x*) = *u*_*s*_(*x, θ*(*x*)) of 𝒜 = 𝒜 (*x, θ*(*x*)), i.e.

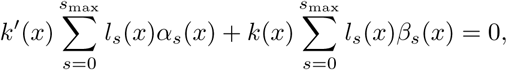

Integrating the above linear homogeneous first-order equation in *k*(*x*) yields

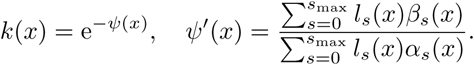

The function *ψ*(*x*) is determined by numerically integrating *ψ*′(*x*). The dependence of the prefactor *k*(*x*) on the protein concentration *x* is exemplified in Figure 6, right panel.

## 7 Discussion

We formulated and investigated a stochastic model for the production of a protein with delayed positive feedback. In the model, the protein is produced in bursts of multiple molecule copies. Newly produced protein molecules are inactive, and become activated by passing through a single activation step; biologically, the step can represent chemical modification, compartmental transport, or other scenarios. Active protein molecules regulate the frequency of bursty production of inactive protein. Such feedback can biologically be realised through transcriptional regulation.

The model incorporates an upper bound *s*_max_ on the number of active protein. If *s*_max_ active protein are already present, a new activation is allowed to occur, but is immediately followed by the removal of the activated molecule; consequently, the number of active protein molecules never exceeds *s*_max_. Thanks to the introduction of the upper bound, a number of crucial steps in the mathematical analysis involve finite, rather than infinite, calculation (e.g. the averaging (5) or the matrix (30)). Without an explicit upper bound in the model, each of these calculations would require an ad-hoc truncation; the explicit inclusion of the upper bound in the model guarantees a consistent use of truncation throughout the entire analysis. In the presented numerical examples, we choose *s*_max_ large enough in order that the results be close to those expected without an upper bound.

We focused on examining the model behaviour in the regime of slow activation. The regime is characterised by activation rates of *O*(1) and production/decay rates of *O*(1*/ε*), where *ε* ≪ 1. Consequently, the inactive protein is present at *O*(1*/ε*) large copy numbers and fluctuates on a *O*(1) slow timescale, whereas the active protein is present at *O*(1) moderate copy numbers and fluctuates on a *O*(*ε*) fast timescale. Performing a quasi-steady-state (QSS) elimination of the fast active protein, we found that the QSS distribution of the active protein is a (truncated) Poisson, and that the inactive protein evolves on the *O*(1) timescale according to a deterministic rate equation (8).

We were specifically interested in situations where the rate equation has two stable steady states. Bistability occurs if the effective feedback response function, which quantifies the feedback of the inactive protein on itself, is sufficiently sigmoid so as to support multiple fixed points. The effective response function is equal to the average value, weighted by the QSS distribution of the active protein, of the original, active-protein-dependent, production rate. As a result of averaging by the noisy active protein, the effective response function is a smoothed-out (or “mollified”) version of the original response function; the requirement that the mollified function be sigmoid implies that the original function must be yet steeper. For simplicity, we used an (infinitely steep) step function in the examples of this paper. Biologically, highly sigmoid feedback responses can be maintained through cooperative binding of the protein to the regulatory DNA sequences.

If the model operates in the slow activation regime and if the limiting rate equation is monostable, then the steady-state distribution of the inactive (active) protein is nearly Gaussian (Poisson); the location of the Gaussian/Poisson mode is dictated by the unique fixed point of the rate equation. If the rate equation is bistable, the distribution of the inactive protein is approximated by a mixture of two small-noise Gaussians, and that of the active protein by a mixture of two (moderate-noise) Poissons; the locations of the Gaussian/Poissonian modes are dictated by the fixed points of the rate equation. In order to obtain asymptotic approximations of the weights of the two modes, one needs to consult (and calculate) a WKB solution to the master equation; doing so was the concern of the bulk of the mathematical analysis presented in this paper. The asymptotic solution agrees well with a numerical solution to the master equation.

The principal step in the calculation of the asymptotic WKB solution is the determination of the WKB potential. The derivative of the potential is formed by the nontrivial heteroclinic connections between the steady states of a Hamiltonian system (Figure 6, full red line). The trivial heteroclinic connections that lie on the *x*-axis (Figure 6, dashed red line) satisfy the rate equation (the one previously established by the QSS analysis). The potential derivative and the deterministic rate have opposite signs: the potential has local minima/maxima where the rate equations has stable/unstable steady states; in other words, the WKB potential is the deterministic rate equation’s Lyapunov function.

Our asymptotic analysis stands on the shoulders of previous analyses (see Introduction for a limited review), and one in particular: Newby and Chapman (2014) study a stochastic gene expression model which is based on different biological assumptions than ours; the commonality is that it features two components with a similar pattern of time and abundance scales. The model of Newby and Chapman (2014) consists of an “internal” finite-state Markov chain (representing promoter states) coupled with an “external” birth and death process (representing protein). The coupling of the two components is in the dependence of transition rates for either component on the current state of both. The deterministic limit in protein dynamics is obtained by reducing both the internal and the external noise. The internal noise is reduced by speeding up the promoter transitions; the external noise is reduced by increasing the protein abundance. If both noise sources are reduced proportionally to each other (and to a small parameter *ε*), the same configuration of time and abundance scales is achieved as in our model in the slow activation regime: the promoter state fluctuates at *O*(1) numbers on a *O*(*ε*) time scale and the protein at *O*(1*/ε*) numbers on a *O*(1) time scale. Like we did here for our model, Newby and Chapman (2014) used the WKB method to describe the large-time behaviour of their model in the *ε* ≪ 1 regime. There are some similarities, as well as differences, between the two models as well as in the methodologies of this paper and Newby and Chapman (2014). Our model features bursting and a stoichiometric connection between the two species that the model of Newby and Chapman (2014) does not. The WKB potential is determined, in both studies, from the condition that a matrix, here (30), be singular. Our matrix is large and sparse, whereas that of Newby and Chapman (2014) is dense and typically small (few gene states). Methodologically, we define the Hamiltonian as the principal eigenvalue (one with the largest real part), where Newby and Chapman (2014) use the determinant, of the matrix. The method of determining the prefactor (Section 6.3) from the higher-order terms of the master equation is the same as used in Newby and Chapman (2014).

In future work, we would like to look beyond the steady state and quantify the transition rates between the modes *x*_−_ and *x*_+_ of the mixture distributions identified in the present paper. It is expected that these rates are proportional to exp(−(*Φ*(*x*_0_) − *Φ*(*x*_±_))*/ε*), i.e. exponentially small as *ε* → 0. The proportionality constant will be determined by matching the WKB solution to a solution of a Fokker–Planck equation in the neighbourhood of the unstable steady state *x*_0_ (Hinch and Chapman, 2005; Bressloff, 2014). We would also like to see the current framework extended to more general distributed delays; we expect that delays composed of *m* slow memoryless steps will lead to a Hamiltonian system in the 2*m*-dimensional Euclidean space.

In summary, we performed a detailed analysis of the steady-state distribution for an autoregulating protein with a large one-step production delay. Previous studies show that deterministically monostable positive feedbacks can exhibit bimodal distributions when formulated stochastically (Singh, 2012; Bokes and Singh, 2019). Our analysis shows that while both monostable and bistable feedbacks can exhibit bimodality at the single-cell level without any time delay, they will converge to different distributions with the inclusion of large delays, and hence provides a novel method to probe the structure of positive genetic feedback circuits.

## Appendix A Linearisation of the Ham. system

Here we derive the expression (50) for the second *θ*-derivative of the Hamiltonian (36). By doing so, we complete the linearisation analysis of the Hamiltonian system (32).

Differentiating ***Av*** = *H****v*** with respect to *θ* twice yields

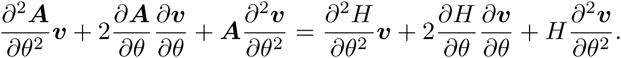

At a saddle with coordinates *x* = *x*_*_ ∈ {*x*_−_, *x*_0_, *x*_+_} and *θ* = 0, we have 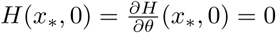 and ***v***(*x*_*_, 0) = ***ρ***(*x*_*_), so that

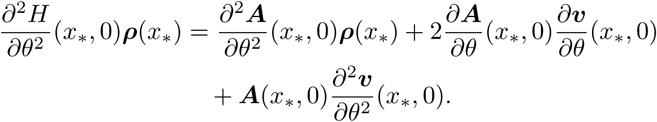

We multiply the above equation by ***u***^**T**^(*x*_*_, 0) = **1**^**T**^ from the left; noting that **1**^**T**^***ρ***(*x*_*_) = 1 and **1**^**T**^***A***(*x*_*_, 0) = **0**^**T**^, we obtain

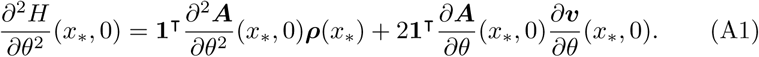

The second derivative of ***A***(*x, θ*) is the matrix representation of the second derivative

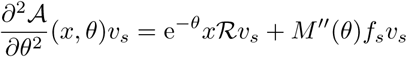

of the operator 𝒜 (*x, θ*) (29). Therefore, the first term on the right-hand side of (A1) satisfies

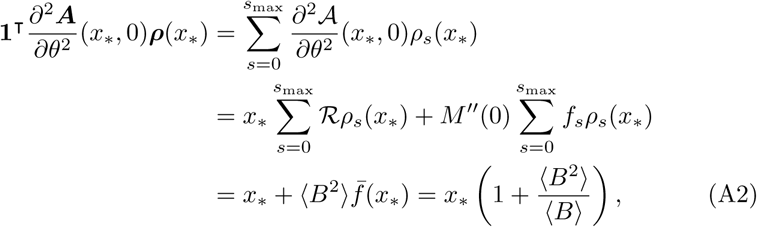

in which we utilised the fact that *x*_*_ is a fixed point of 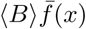.

The second term on the right-hand side of (A1) involves the derivative of the principal eigenvector with respect to *θ*. Differentiating ***Av*** = *H****v*** with respect to *θ* and inserting *x* = *x*_*_ and *θ* = 0 yields

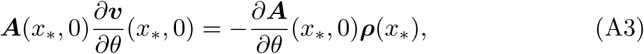

Solving the inhomogeneous linear algebraic system (A3) yields

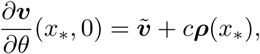

where *c****ρ***(*x*_*_) is a representant of the (right) kernel of ***A***(*x*_*_, 0) and

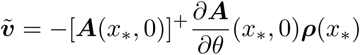

is a least-squares solution to (A3), with ***A***^+^ denoting the pseudo-inverse of ***A***.

The second term in (A1) therefore satisfies

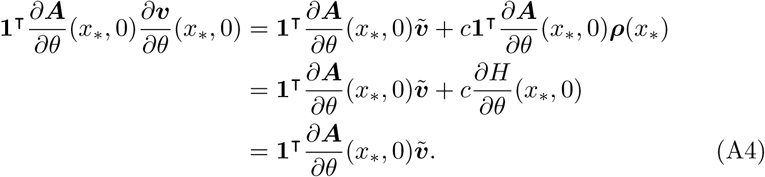

Inserting (A2) and (A4) into (A1) recovers (50).

